# Disrupted Emotional Neural Synchrony in Schizophrenia Revealed by Intersubject Correlation of Naturalistic fMRI

**DOI:** 10.64898/2026.04.13.718247

**Authors:** Carla Pallavicini, Elsa Yolanda Costanzo, Laura Alethia de la Fuente, Mariana Nair Castro, Salvador Martín Guinjoan, Enzo Tagliazucchi, Mirta Villarreal

## Abstract

**Background:** Schizophrenia is marked by impairments in emotional processing and social cognition, yet traditional neuroimaging paradigms often lack the ecological validity to capture these deficits in real-world contexts.

**Methods:** In this study, we used intersubject correlation (ISC) analysis of functional MRI data to examine shared neural representations of naturalistic visual narratives in individuals with schizophrenia and healthy controls. Participants viewed short films designed to evoke happy, sad, and emotionally neutral responses, allowing us to compare how synchronized brain activity varied with emotional content across and within groups.

**Results:** Healthy controls showed greater ISC in regions associated with affective salience, emotion recognition, and social understanding, including the amygdala, insula, and temporal cortices. In contrast, participants with schizophrenia displayed higher synchrony in visual, subcortical, and frontal areas, suggesting a reliance on perceptual and executive systems. To isolate the effects of emotion from general visual processing, we compared ISC during emotional clips relative to neutral videos. This revealed significantly reduced synchrony in the bilateral amygdala in patients, highlighting a core dysfunction in affective engagement. Interestingly, neutral stimuli elicited unexpectedly strong synchronization in frontal and limbic regions in the schizophrenia group, possibly reflecting altered salience attribution to ambiguous or emotionally ambiguous content.

**Conclusions:** These results point to a functional reorganization of affective processing in schizophrenia, where impaired limbic recruitment is accompanied by compensatory engagement of perceptual and cognitive control networks. ISC during naturalistic stimulation emerges as a powerful tool for capturing subtle disruptions in shared emotional experience in psychiatric populations.

## Introduction

Schizophrenia is a severe and chronic psychiatric disorder that affects approximately 24 million people worldwide, or about 1 in 300 people (0.32%) globally, and 1 in 222 adults (0.45%) according to the World Health Organization (2022) [1,2]. It typically manifests in late adolescence or early adulthood and can cause substantial impairments in perception, cognition, emotion, and social functioning [3, 4]. The disorder is associated with symptoms such as hallucinations, delusions, disorganized thinking, and cognitive deficits, making it one of the leading causes of disability worldwide [4, 5, 6, 7, 8, 9, 10, 11]. Despite decades of research, the underlying neural mechanisms of schizophrenia remain only partially understood, particularly in how patients process complex emotional and social information [12].

A growing body of literature has documented that schizophrenia is characterized by fundamental disruptions in emotional processing and social cognition [10, 13, 14, 15]. Patients often show blunted affect [16], misinterpretation of emotional cues [17], and difficulties in empathy and theory of mind [18, 19, 20, 21]. A recent meta-analysis by Jáni & Kašpárek [10] confirmed that schizophrenia patients exhibit decreased activation in the amygdala, thalamus, insula, anterior cingulate cortex (ACC), and medial prefrontal cortex (mPFC) during emotion recognition tasks, while also displaying increased activation in parietal and motor-related regions, possibly reflecting compensation. These abnormalities suggest that emotional information in schizophrenia may be processed in a more perceptually driven and less socially integrated manner, with additional reliance on regions outside the classical emotion recognition network. Furthermore, theory of mind impairments were associated with reduced recruitment of the mPFC and right insula, areas typically involved in understanding others’ mental states [10, 22, 23]. These neural patterns suggest that emotional and social cognitive deficits in schizophrenia may stem from both underactivation of core socio-emotional systems and atypical reliance on compensatory perceptual strategies.

Recent advances in cognitive neuroscience have emphasized the use of naturalistic stimuli, such as movies, stories, or real-life narratives, to more faithfully engage brain systems under conditions resembling everyday experiences [24, 25, 26]. These stimuli are continuous and multidimensional, embedding visual cues in multi-object scenes and involving complex motion. They permit free eye movement, allowing visual processing to dynamically interact with context, emotional valence, and other sensory modalities [26, 27, 28]. Importantly, they engage conscious perception in a natural and temporally extended way. However, due to their complexity, naturalistic stimuli cannot be analyzed using traditional task-based functional magnetic resonance imaging (fMRI) contrasts, which assume discrete events or conditions. This is particularly relevant in schizophrenia, where disruptions in context integration [29,30], temporal binding [31, 32], and attentional allocation [33,34, 35] may be more evident during naturalistic scenarios than in block or event-related paradigms.

To address these analytical challenges, researchers have developed methods like Intersubject Correlation (ISC) analysis [36]. ISC measures the similarity of brain responses across individuals exposed to the same naturalistic stimulus, without requiring an explicit model of the stimulus [24, 26, 37, 38, 39, 40]. This approach captures temporal neural alignment across individuals, offering a powerful window into shared perceptual and cognitive processes. In clinical populations such as schizophrenia, where individual variability is high and task compliance can be difficult, ISC offers an elegant framework to assess collective neural engagement with emotionally rich content.

In this fMRI study, we investigated how age- and gender-matched individuals with schizophrenia and healthy controls respond to naturalistic audiovisual narratives with emotional content. We presented 34 participants (20 controls and 14 individuals diagnosed with schizophrenia; Table 1) with film segments eliciting happy, sad, and neutral emotional responses. Using ISC analysis, we examined the extent to which patients and controls exhibit synchronized neural activity in response to emotional stimuli, and whether schizophrenia is associated with alterations in intersubject neural coherence. This approach revealed distinct patterns of neural synchrony between patients and controls, offering new insights into how emotional content is processed and shared in schizophrenia.

**Table 1.** Demographic characteristics.

|  | **Patients** | **Controles** | **Statistics** | **P** |
| --- | --- | --- | --- | --- |
|  | **(n = 14)** | **(n=20)** |  |  |
| Gender (male/female) | 9/5 | 9/11 | $\chi^2 = 1.229$ | 0.315 |
| Age (years) | 30.3 ± 6.4 | 25.9 ± 5.5 | t = 2.173 | 0.035 |
| Education (years) | 12.6 ± 2.2 | 17.3 ± 3.2 | t = -4.768 | < 0.05 |
| HAM-D Baseline | 10.0 ± 8.6 | 2.0 ± 2.2 | t = 3.978 | < 0.05 |
| WAT | 33.5 ± 5.8 | 35.6 ± 4.1 | t = -1.268 | 0.214 |
| PANSS total score | 80.6 ± 33.1 | - | - | - |
| Positive Subscale | 17.3 ± 8.0 | - | - | - |
| Negative Subscale | 27.5 ± 11.1 | - | - | - |

Shown are mean ± SD of each variable, p value for independent samples t test (numeric variables) or chi-square test (categorical variables). HAM-D: Hamilton Depression Scale- WAT: Word Accentuation Test. PANSS: Positive and Negative Symptoms Scale.

## Methods

### Participants

Thirty-four participants, 14 psychiatry outpatients with schizophrenia (SCH, 5 females, aged 30.3 ± 6.4 years) from the Psychiatry Service at Fleni and 20 healthy controls (HC, 11 females, aged 25.9 ± 5.6 years), were recruited. Patients were invited to participate in the study if they (a) received a DSM-IV-TR (American Psychiatric Association, 1994) diagnosis of schizophrenia (any subtype), (b) were aged 18–65 years, and (c) had been on the same medications for at least two weeks. In terms of pharmacological treatment, 58% of patients were receiving antipsychotics, 24% antidepressants, and 24% anxiolytics^1^. General exclusion criteria included: (a) present or past serious medical illness, including neuropsychiatric disorders (other than schizophrenia for SCH), (b) misuse or addiction to illegal substances in the previous 6 months, and (c) contraindications for magnetic resonance imaging (e.g., metal implants, pacemakers, claustrophobia). Additional exclusion criteria for healthy controls included any lifetime DSM-IV-TR Axis I diagnosis (as assessed by a consultant psychiatrist) and any history of treatment with antidepressants, antipsychotics, or mood stabilizers. All individuals provided informed consent as approved by the local bioethics committee: Comité de Ética en Investigación FLENI. The study was conducted in accordance with the ethical standards set by the 1964 Declaration of Helsinki.

### Screening tests

All participants were screened for premorbid intelligence with the Word Accentuation Test (WAT) [41, 42] and for depressive symptoms with the Hamilton depression test (HAM-D) [43]. All patients were evaluated using Positive and Negative Symptoms Scale (PANSS) [44] to measure psychotic symptom severity.

### Stimuli and Experimental Design

Subjects participated in an affective visual procedure, consisting of the presentation of video clips of one minute duration in two different affective conditions (happiness and sadness), plus a control condition (neutral), see Figure 1. Participants were told in Spanish: “During this task, I would like you to try to become happy [or sad], helping yourself with the videos showing such emotion”.

**Figure 1.**
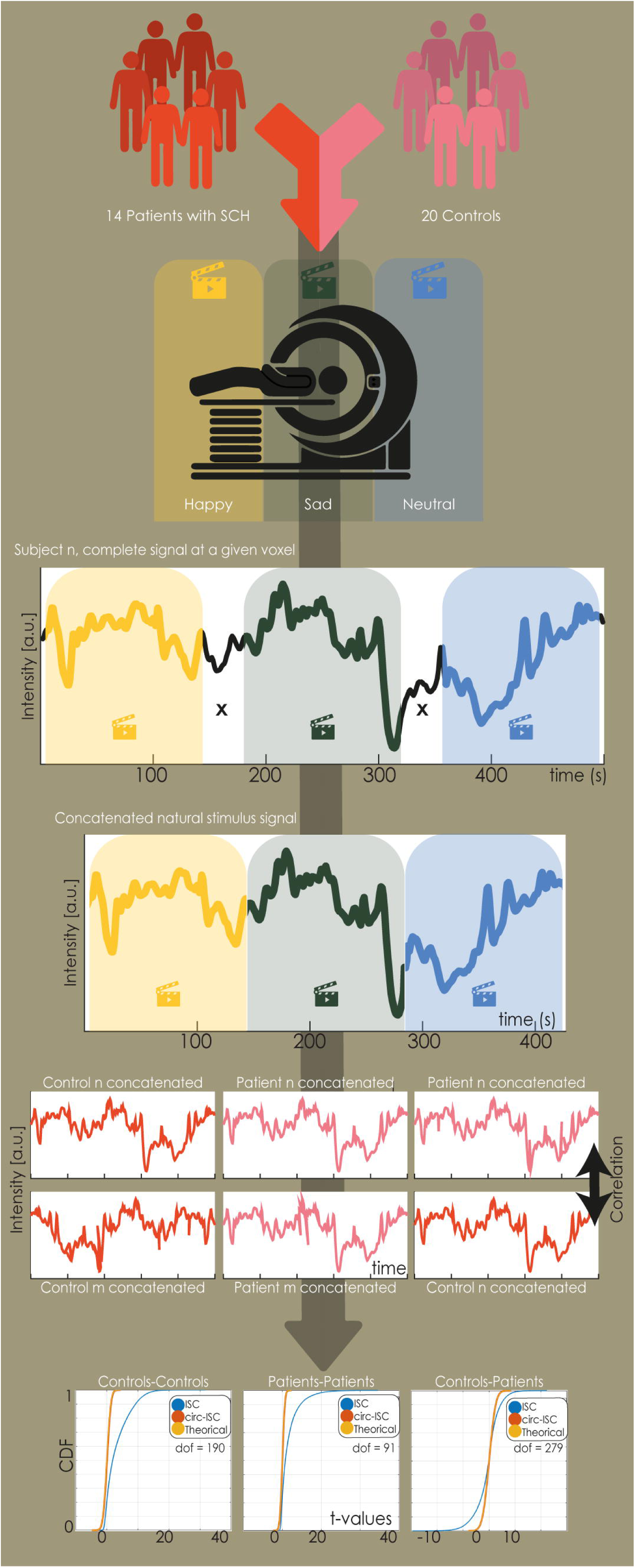
Overview of Experimental Paradigm and Analysis Pipeline. Schematic representation of the experimental design and analysis procedure. Participants (schizophrenia and control groups) watched a naturalistic emotional film while undergoing fMRI. After preprocessing, voxel-wise inter-subject correlation (ISC) was computed within each group. Group-average ISC maps were generated and contrasted between groups. Statistical significance was assessed using nonparametric permutation testing and corrected for multiple comparisons. This pipeline enabled identification of brain regions with reduced or increased synchronization in schizophrenia during emotional movie viewing.

The experimental design consisted of 3 sessions: one for sad responses, another for happy responses, and a third for neutral control. Each session consisted of 3 blocks of task condition (sad [happy] [neutral] videos) alternated with 3 blocks of baseline (fixation cross) of 60 s each block. The blocks were presented randomly to each participant.

### fMRI Data Acquisition

MRI data were acquired on a 3 T GE HDx scanner with an 8 channel head coil. Change in blood-oxygenation–level-dependent (BOLD) T2* signal was measured using a gradient echo-planar imaging (EPI) sequence. Thirty contiguous slices were obtained in the AC-PC plane (TR: 2 s, TE: 30 ms, flip angle: 90◦, FOV: 24 cm, 64 × 64 pixels per inch matrix, voxel size = 3.75 × 3.75 × 4). A structural MRI was acquired with the T1-weighted 3D fast SPGR-IR sequence (120 slices, 1.2-mm thick slices, TR = 6.604 ms, TE = 2.796 ms, flip angle 15◦, FOV 24 cm, 256 × 256 matrix).

### Preprocessing

For each participant and for each brain state, we used FSL tools to extract and average the BOLD signals from all voxels. Standard preprocessing steps included brain extraction (BET), spatial smoothing with a 5 mm full-width at half maximum (FWHM) Gaussian kernel, and bandpass temporal filtering in the 0.01–0.1 Hz range. Functional images were then registered to standard space (2 mm MNI template) and resampled to a final voxel size of 4 mm isotropic, yielding images of dimensions 45 × 54 × 45.

### Intersubject Correlation (ISC)

The ISC approach in this study was adapted from the methodology described by Hasson et al. (2004) [36]. Each participant’s preprocessed functional data were transformed to MNI space and segmented according to the corresponding narrative condition. Frames corresponding to baseline fixation periods were excluded from the analysis to focus exclusively on stimulus-driven brain responses. Furthermore, the fMRI time series corresponding to the viewing of each video clip within the same emotional valence category (happy, sad, or neutral) were concatenated, creating a single continuous dataset per emotion for each participant. A general linear model (GLM) was applied for each narrative, incorporating temporal drift regressors modeled with cubic splines (one per two minutes of scanning), six motion parameters, and a whole-brain average signal. These nuisance regressors were not convolved with a hemodynamic response function, and the first 6 seconds of each narrative were excluded to minimize onset-related transients. Residual time series from the GLM were retained for ISC computation.

Pairwise ISC maps were computed for all subject pairs within each group using a custom MATLAB script, resulting in 91 pairs for the schizophrenia group and 190 pairs for the controls. The Pearson correlation coefficient (r) between residual time courses at each voxel was calculated and subsequently transformed using Fisher’s z transformation to normalize the distribution (Figure 1)

To isolate emotional processing from baseline sensory responses, we implemented neutral-subtracted contrasts for both the schizophrenia and control groups. Specifically, for each voxel, we computed ISC values during emotionally valenced videos (happy or sad) and subtracted ISC values during neutral videos, resulting in a voxel-wise emotional-resonance index. This subtraction helps eliminate shared variance attributable to general engagement with the video format.

### Group analyses

Group analyses were performed to determine where ISC significantly deviated from zero, and where the mean of the ISC distributions differed between patients and controls. Given the statistical dependence between the 91 and 190 pairs of ISC maps, we first implemented a null model to estimate the distribution of the t statistic, following the analysis implemented in Wilson et al. [37]. For this purpose, we temporally shifted each subject’s time series by a random number of volumes, effectively breaking temporal alignment while maintaining individual autocorrelation. This generated synthetic datasets where any observed correlation would be due to chance. The resulting null distributions closely approximated the theoretical distribution of uncorrelated data with equivalent degrees of freedom, corresponding to 91 for the schizophrenia group, 190 for the controls, and 279 for the between-group comparison (Supplementary Figure 1). Statistical parametric maps for each group and the between-group comparison were thresholded at p<0.001, with an additional threshold of p<0.05, cluster-level, with Family-Wise Error (FWE) correction as implemented in SPM12.

## Results

### Demographic characteristics

Table 1 shows demographic and clinical characteristics of participants. Both groups were similar regarding gender and depressive symptoms. Patients with schizophrenia were older and had fewer years of education than control subjects (p < 0.05; Table 1).

### Intersubject Correlation Within Groups

We first examined brain regions showing significant intersubject correlations within each group across the three emotional conditions: happy, sad, and neutral. The ISC analysis revealed distinct as well as overlapping patterns of synchrony across healthy controls and patients. Significant correlated areas are illustrated in Figure 2 and summarized in Supplementary Table 1.

**Figure 2.**
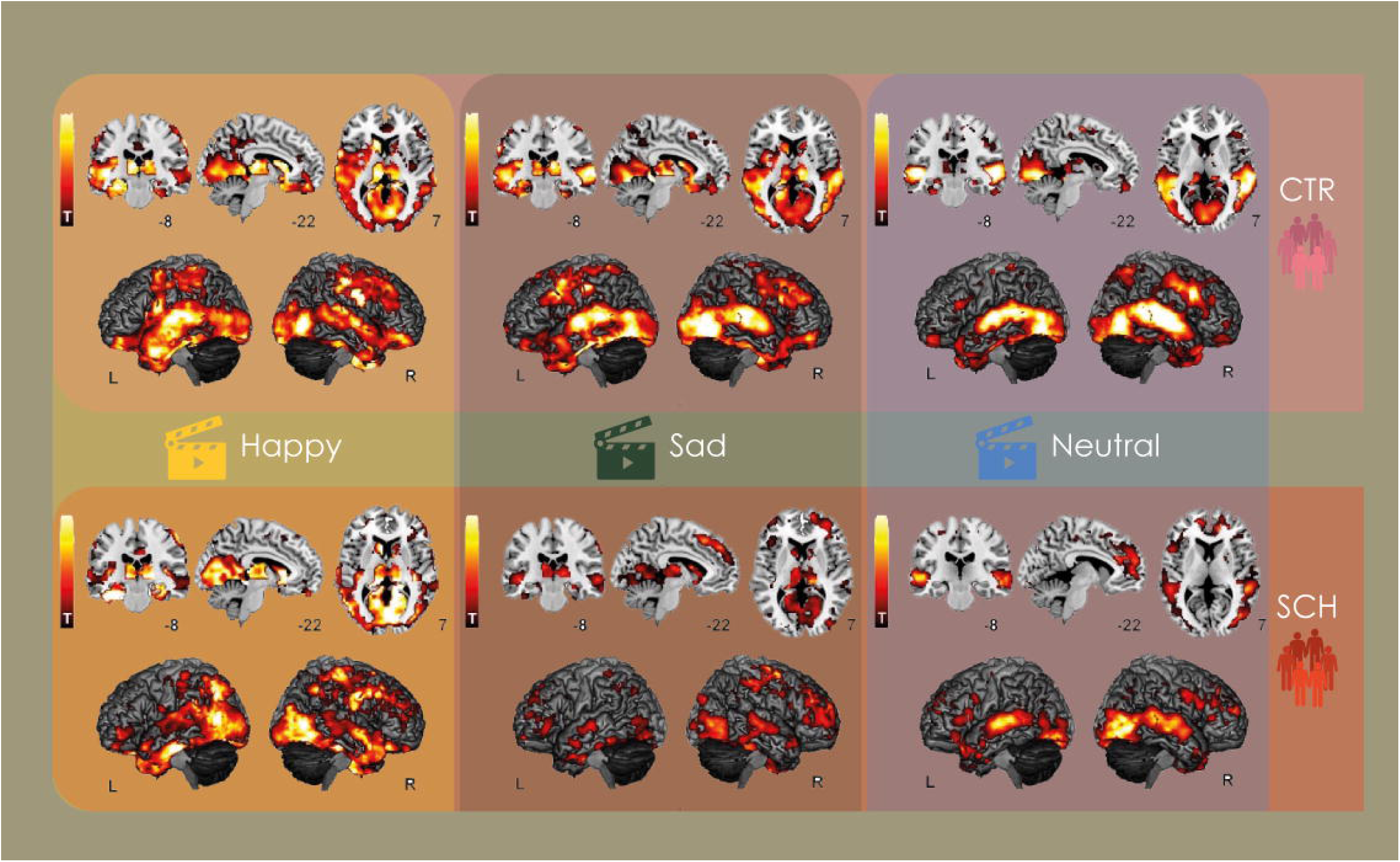
Group Differences in Inter-Subject Correlation During Naturalistic Movie Viewing. Brain regions showing significantly reduced ISC in patients with schizophrenia (orange) and regions showing significantly higher ISC in patients compared to controls (pink). Statistical maps were corrected for multiple comparisons using permutation testing and displayed at cluster-wise corrected p < 0.05. Reduced ISC was predominantly observed in visual and limbic areas, while increased ISC appeared in frontal and temporal regions associated with higher-order processing. ISC calculated over happy (yellow), sad (green) and neutral (blue) videos.

During the happy condition (Figure 2, left panels), both groups exhibited significant ISC in early visual areas such as the *calcarine sulcus* and *lingual gyrus*, indicating shared low-level visual processing. Controls showed broader ISC patterns encompassing higher-order temporal regions including the *middle temporal gyrus*, *superior temporal gyrus*, and *inferior temporal gyrus*, which are associated with audiovisual integration and emotional content processing. Notably, the *caudate* also showed synchronization, likely reflecting engagement in reward-related processing. In contrast, patients showed less extensive ISC in temporal regions but exhibited additional ISC in the *middle occipital gyrus* and *thalamus*, areas involved in sensory relay and visual processing, possibly reflecting a more bottom-up processing bias.

In the sad condition (Figure 2, middle panels), both groups again showed shared ISC in the *lingual gyrus* and *middle temporal gyrus*. Controls demonstrated ISC in regions typically involved in affective processing such as the *amygdala*, *fusiform gyrus*, and *precuneus*, the latter known for its role in self-referential and emotional introspection. Patients, by contrast, recruited the *parahippocampal gyrus*, *calcarine sulcus*, *thalamus*, and *inferior temporal gyrus*, with a notable absence of the amygdala, suggesting altered engagement with emotionally salient content.

For the neutral stimuli (Figure 2, left panels), shared ISC across both groups was observed in visual processing regions such as the *inferior occipital gyrus*, *middle temporal gyrus*, *inferior temporal gyrus*, and *lingual gyrus*. Controls also recruited the *fusiform gyrus* and *superior temporal gyrus*, involved in face and voice processing. Patients, on the other hand, exhibited additional ISC in the *superior frontal gyrus*, a region associated with self-monitoring and attentional control, which may reflect compensatory mechanisms or increased cognitive effort in decoding neutral narratives.

### Emotion-Modulated ISC Within Groups

Next, we explored within-group ISC differences to assess how emotional content modulated intersubject synchrony in each group, illustrated in Figure 2 and Supplementary Figure 1. Comparisons were made between the happy and neutral, and sad and neutral conditions, with statistical maps thresholded at *p* < 0.005 uncorrected and *p* < 0.05 cluster-level corrected. Results are summarized in Supplementary Table 2.

In the happy vs. neutral contrast (Figure 3, left), healthy controls (top panel) showed increased ISC in the *insula*, *hippocampus*, and *amygdala*, all regions deeply involved in emotional salience, memory encoding, and affective valuation. Additional ISC emerged in the *inferior temporal gyrus*, while decreased ISC was observed in the *superior temporal gyrus* and *supplementary motor area*, potentially reflecting attenuated auditory or motor mirroring engagement under heightened emotional content. In contrast, schizophrenia patients (bottom panel) demonstrated increased ISC in the *calcarine sulcus*, *lingual gyrus*, *caudate*, *thalamus*, *middle temporal gyrus*, and *postcentral gyrus*, regions linked to visual, sensory-motor, and relay processing, again suggesting a bias toward perceptual processing pathways.

**Figure 3.**
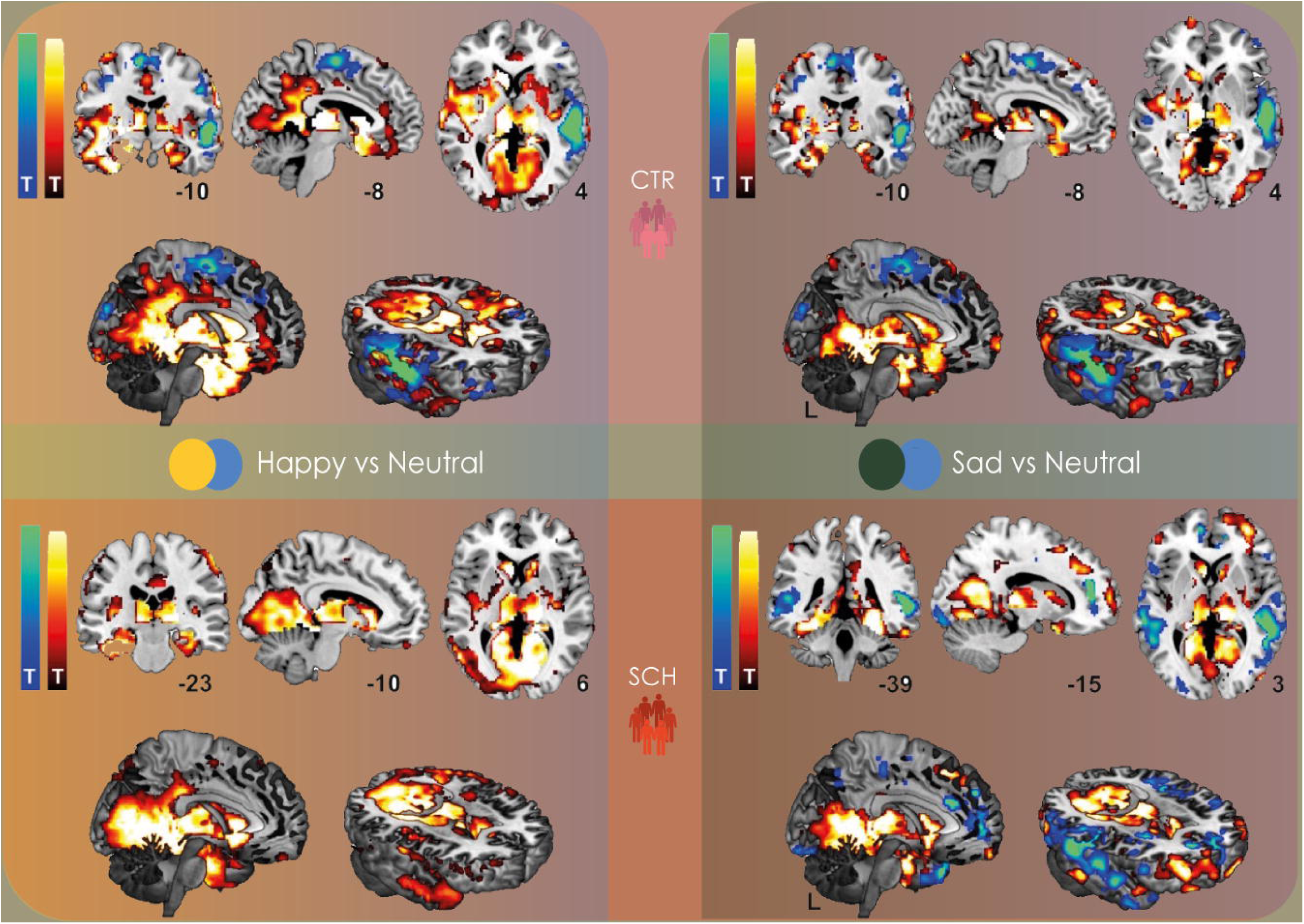
Emotional Context Modulates ISC Within Groups. Within-group ISC contrasts comparing emotional (happy or sad) versus neutral video segments in healthy controls and patients with schizophrenia. Warm colors indicate regions with higher ISC during emotional compared to neutral scenes. Results are shown at p < 0.005 uncorrected and p < 0.05 cluster-level corrected. Both groups showed increased ISC in response to emotional content, though with partially distinct topographies.

The sad vs. neutral contrast is shown in the right panels of Figure 3, controls (top section) again exhibited increased ISC in the *insula*, *lingual gyrus*, *thalamus*, and *amygdala*, consistent with emotional integration and affective arousal. However, decreased ISC was found in the *superior temporal gyrus* and *middle temporal gyrus*, suggesting that emotionally negative stimuli may suppress shared responses in these auditory-social processing regions. Patients (bottom section) showed increased ISC in the *lingual gyrus*, *fusiform gyrus*, and *thalamus*, with additional decreased ISC in the *anterior cingulate cortex* and *middle temporal gyrus*. This pattern reflects partial engagement of emotional-perceptual networks, alongside reduced coherence in frontal integrative hubs.

### Between-Group Differences in ISC Across Emotional Conditions

To further characterize group differences in intersubject neural synchrony, we compared ISC values between controls and patients with schizophrenia for each emotional condition individually, as well as for happy and sad conditions relative to the neutral baseline, as shown in Figure 4. Statistical thresholds were set at *p* < 0.005 uncorrected and *p* < 0.05 corrected at the cluster level.

**Figure 4:**
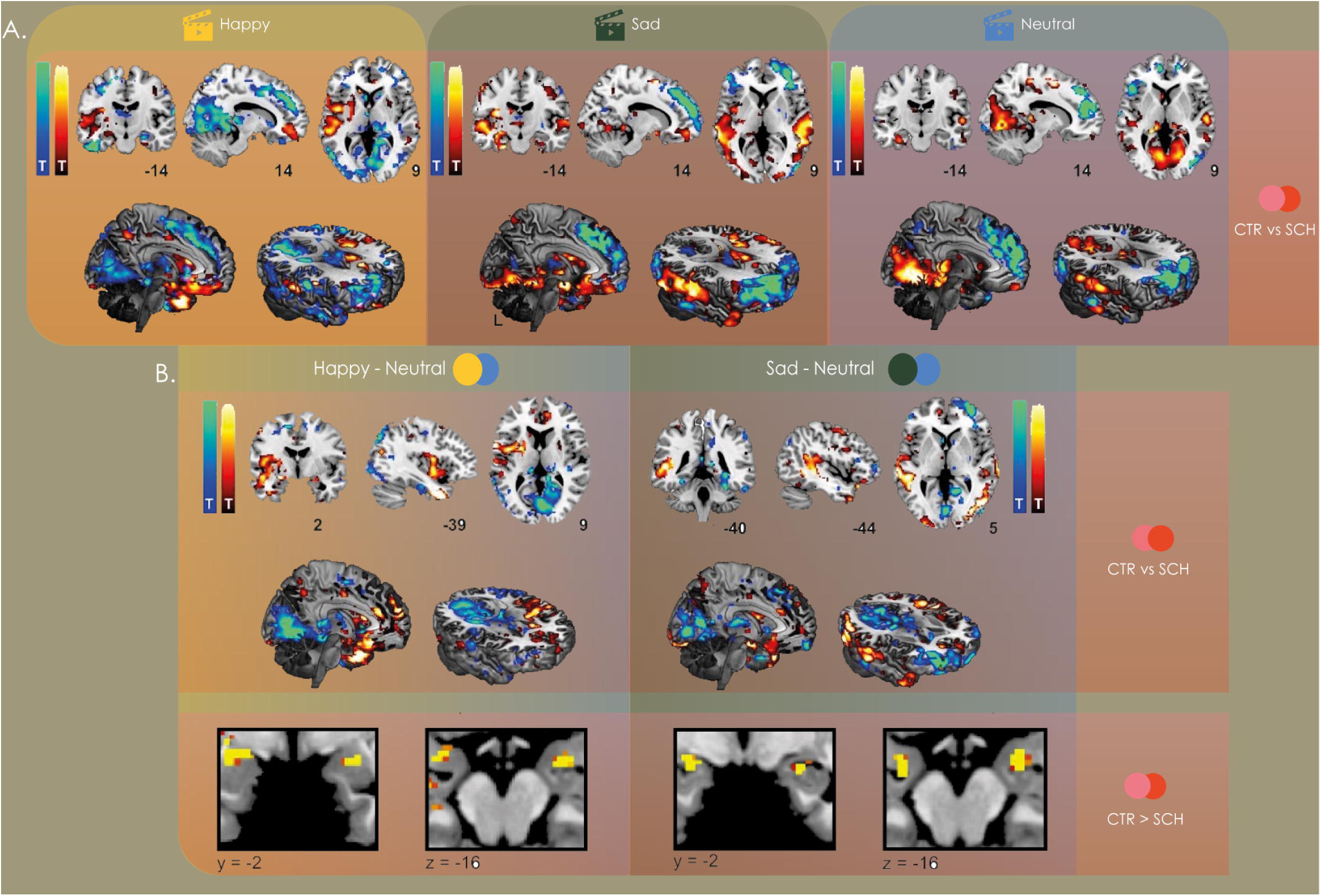
Reduced Inter-Subject Correlation in Emotion-Processing Regions in Schizophrenia. A) Between-group differences in ISC for the comparison of controls vs. schizophrenia and the three conditions. B) Top: Differences between controls and schizophrenia for the conditions happy and sad with neutral subtracted as a baseline (thresholded at p < 0.005 uncorrected and p < 0.05 corrected at the cluster level). Bottom: Detail of the bilateral amygdala showing decreased ISC for schizophrenia vs. controls in the conditions happy – neutral and sad – neutral (thresholded at t > 4).

In the happy condition (Figure 4A, left), controls exhibited significantly greater ISC than schizophrenia patients in regions related to emotional and social processing, including the *insula*, *middle temporal gyrus*, *precentral gyrus*, *amygdala*, and *inferior temporal gyrus* (Figure 4A, left). These areas are implicated in affective salience, motor mirroring, and audiovisual integration. Conversely, schizophrenia patients showed greater ISC compared to healthy controls in the *superior frontal gyrus*, *orbitofrontal cortex*, *fusiform gyrus*, *angular gyrus*, and in regions encompassing parts of both the *middle temporal* and *precentral gyri*. These areas are often associated with attentional control, self-referential processing, and compensatory cognitive strategies.

For the sad condition (Figure 4A, middle), similar patterns emerged. Controls showed stronger ISC in the *insula*, *superior temporal gyrus*, *middle temporal gyrus*, *precentral gyrus*, *amygdala*, and *fusiform gyrus*, regions typically engaged during emotional perception and social cognition. In contrast, schizophrenia patients showed higher ISC in the *supramarginal gyrus*, *superior frontal gyrus*, *inferior temporal gyrus*, *middle frontal gyrus*, *middle occipital gyrus*, and *supplementary motor area*, a pattern suggestive of greater reliance on perceptual and motor planning areas, with reduced involvement of limbic-affective networks (Figure 4A, middle).

In the neutral condition (Figure 4A, right), controls showed higher ISC in the *lingual gyrus*, *superior temporal gyrus*, *calcarine sulcus*, *middle temporal gyrus*, *thalamus*, and *parahippocampal gyrus*, consistent with coordinated visual and memory-related processing. Schizophrenia patients exhibited stronger ISC in frontal midline and regulatory regions, including the *superior frontal gyrus*, *inferior frontal gyrus*, *anterior cingulate cortex*, and *orbitofrontal cortex*, areas often implicated in compensatory engagement or increased internal monitoring demands (Figure 4A, right).

To further clarify the role of emotional response, we compared happy and sad conditions against the neutral baseline across groups.

In the happy vs. neutral contrast, controls demonstrated stronger ISC in key affective and integrative regions including the *insula*, *inferior temporal gyrus*, *amygdala*, *middle temporal gyrus*, *superior frontal gyrus*, and *putamen*. These regions are involved in emotional processing, social attention, and motivational salience. Schizophrenia patients, however, showed higher ISC in posterior and subcortical regions such as the *parahippocampal gyrus*, *lingual gyrus*, *inferior occipital gyrus*, *calcarine sulcus*, *thalamus*, and *cuneus*, suggesting a visual-perceptual shift and diminished limbic synchrony (Figure 4B, top left).

For the sad vs. neutral contrast, healthy controls again showed elevated ISC in emotional hubs like the *insula*, *amygdala* (notably with repeated clusters), *middle temporal gyrus*, *fusiform gyrus*, and *inferior frontal cortex*. Schizophrenia patients exhibited increased ISC in the *calcarine sulcus*, *middle frontal gyrus*, *thalamus*, *middle occipital gyrus*, *superior frontal gyrus*, and *precentral gyrus*. These regions reinforce the interpretation of enhanced reliance on perceptual-motor and dorsal frontal networks in schizophrenia during emotionally charged narratives (Figure 4B, top right).

Details of the bilateral amygdala are presented in the bottom panels of Figure 4B, showing decreased ISC for schizophrenia vs. controls in the conditions happy minus neutral and sad minus neutral (thresholded at t > 4) further supports this interpretation. These results are visualized in Figure 4 and detailed in Supplementary Tables 3 and 4.

## Discussion

This study used intersubject correlation (ISC) analysis to investigate how individuals with schizophrenia and healthy controls synchronize their neural responses when exposed to naturalistic emotional narratives. Our results reveal distinct group-level patterns, with healthy controls showing greater intersubject neural synchrony in regions involved in affective salience, social cognition, and integrative processing. In contrast, schizophrenia patients predominantly exhibited synchrony in perceptual, subcortical, and frontal regions commonly associated with visual processing, attentional control, and executive compensation.

The observed reduction in synchronized activity in canonical emotional networks in schizophrenia aligns with extensive literature on emotion processing deficits in this population [10, 13, 14, 15]. Previous research has consistently reported diminished activity in the amygdala, anterior insula, and medial prefrontal cortex during emotional tasks [10, 45, 46]. Our findings extend this work by showing that these alterations are also reflected in the temporal coordination of neural responses across individuals, highlighting atypical patterns in the processing and sharing of emotional information.

A particularly robust finding in our study was the decreased ISC in the bilateral amygdala in schizophrenia patients compared to controls during both emotionally valenced conditions (happy and sad) when contrasted against the neutral baseline. The amygdala plays a central role in affective salience detection and emotional learning [47], and its diminished involvement in patients provides compelling evidence for a disruption in emotional resonance: the capacity to align affectively with external emotional cues [17, 48, 49]. Crucially, this result emerged even when controlling for shared perceptual input through neutral-subtracted contrasts, indicating that it reflects a core emotional processing deficit rather than a confound driven by basic sensory differences. This interpretation is reinforced by previous fMRI studies reporting reduced amygdala activation in schizophrenia during emotion recognition [46, 48, 49, 50]. Furthermore, other key regions with reduced ISC in our study, including the insula, thalamus, and caudate, have also been repeatedly implicated in dysfunctional emotional and social processing in schizophrenia [10, 23, 51]. Together, these results highlight the value of naturalistic ISC as a tool for capturing distributed neural disruptions in affective processing that are otherwise difficult to detect through traditional task paradigms.

Interestingly, ISC maps for schizophrenia patients frequently involved dorsal and posterior regions, such as the calcarine sulcus, cuneus, thalamus, and middle occipital gyrus, across all emotional conditions. These areas support low-level visual encoding and perceptual integration, suggesting that patients may process complex audiovisual stimuli with an overreliance on bottom-up features, a tendency that could be linked to the presence of positive symptoms, such as hallucinations or aberrant salience attribution. At the same time, enhanced ISC in frontal regions, including the superior frontal gyrus and anterior cingulate cortex, points toward possible compensatory strategies [10]. These areas are often implicated in self-monitoring and cognitive control [22], which may become hyper-recruited when patients struggle to extract or relate to emotional meaning.

These findings align with previous neuroimaging studies demonstrating reduced engagement of key emotion-processing regions in schizophrenia. For instance, Knobloch et al. [46] reported that patients with schizophrenia exhibit decreased activation in the anterior insula, caudate nucleus, thalamus, and inferior frontal gyrus during tasks involving static and dynamic emotional stimuli. Notably, many of these same regions, particularly the anterior insula and thalamus, emerged in our ISC analyses as showing reduced intersubject synchrony in the patient group. More recent work has expanded on these findings, demonstrating that increased amygdala–insula and amygdala–inferior frontal gyrus connectivity during explicit emotional face processing is associated with greater symptom severity and poorer social functioning across psychotic disorders [52]. This convergence highlights the functional importance of these regions in emotional communication and supports our interpretation that schizophrenia involves a breakdown in coordinated emotional processing, even in the absence of overt behavioral deficits. It also highlights the utility of ISC as a sensitive and ecologically valid measure for detecting functional disruptions that may be subtle yet clinically meaningful.

One intriguing finding concerns the “neutral” condition. While ostensibly affectively flat, the neural synchrony patterns in schizophrenia suggest that such stimuli may paradoxically carry more emotional or personal salience for patients than the emotionally labeled clips. This could reflect an altered salience attribution mechanism, a hallmark of schizophrenia [53], where ambiguity or neutrality is perceived as unpredictable or emotionally ambiguous, thereby engaging greater internal resources. The fact that patients displayed marked ISC in frontal and limbic-paralimbic areas under the neutral condition reinforces this hypothesis.

The use of neutral-subtracted contrasts (e.g., happy–neutral, sad–neutral) proved particularly informative [54]. Naturalistic stimuli inherently evoke high perceptual alignment (e.g., all participants visually follow the same film) but the subtraction of neutral ISC helps isolate regions where emotional content specifically drives intersubject synchrony [55]. This approach avoids conflating perceptual entrainment with affective processing and allowed us to capture the unique emotional resonance, or its disruption, in each group.

ISC during naturalistic stimulation emerges from this study as a powerful and noninvasive approach to detect subtle disruptions in shared emotional experience, a hallmark feature of schizophrenia [48]. Unlike traditional cognitive tasks, naturalistic paradigms require minimal instruction, allow for unconstrained attention, and closely resemble real-life emotional situations [55]. This makes ISC particularly suitable for use in clinical and high-risk populations, where task compliance or interpretability may be limited [56]. Because the method relies on the temporal coherence of spontaneous neural responses across individuals, it provides a window into the brain’s capacity to resonate with shared social and emotional signals [36, 37]. Importantly, these neural synchrony signatures may precede the overt onset of psychotic symptoms [57], making ISC a promising candidate for identifying at-risk individuals or subclinical traits. Given its safety, simplicity, and ecological validity, ISC could contribute to early detection efforts, phenotyping variability within schizophrenia, and evaluating treatment effects on emotional engagement and social functioning.

This study has several limitations that warrant consideration. While the overall sample size was modest, ISC inherently capitalizes on pairwise comparisons between participants, which exponentially increases the number of data points used in statistical analyses. As such, the effective sample size for ISC is substantially higher than the number of individual subjects, allowing for robust voxel-wise comparisons even in smaller cohorts [58]. Nonetheless, while similar sample sizes have been reported in previous schizophrenia studies [17, 20, 34, 59, 60, 61], the generalizability of our findings should be confirmed in larger and more diverse cohorts. Additionally, although all patients met diagnostic criteria for schizophrenia, many, as expected, exhibited co-occurring depressive symptoms [6], including some with clinically significant scores, which could contribute to the observed disruptions in affective neural synchrony [10,45]. The possibility that these findings reflect broader affective dysregulation, rather than schizophrenia-specific processes, cannot be ruled out. Moreover, patients were under stable pharmacological treatment, and the heterogeneity of psychotic symptoms, including those involving perceptual disturbances, may have interacted with responses to the naturalistic emotional stimuli in ways that remain difficult to fully disentangle [5, 6, 29, 30, 48, 53]. Future studies should aim to control for symptom severity, mood comorbidities, and medication status to more precisely isolate the neural signatures of emotional processing deficits in schizophrenia.

Altogether, the present study supports a model in which schizophrenia is marked by diminished shared emotional experience at the neural level, possibly underpinned by impaired limbic activation and an overreliance on perceptual and cognitive control systems. The amygdala findings, in particular, provide a striking demonstration of the disorder’s core affective dysfunction. These insights reinforce the utility of naturalistic paradigms combined with ISC analysis as a promising framework to investigate social-emotional impairments in clinical populations.

## Financial Disclosures

Dr. Salvador Guinjoan reports receiving research funding from the National Institutes of Health (NIH) under grants 1R01MH137331-01A1 and 1R21MH138011-01. Dr. Carla Pallavicini reports receiving funding from the European Union’s Marie Skłodowska-Curie Actions (MSCA) program (GradStim – 101064772).

All other authors report no biomedical financial interests or potential conflicts of interest.

---

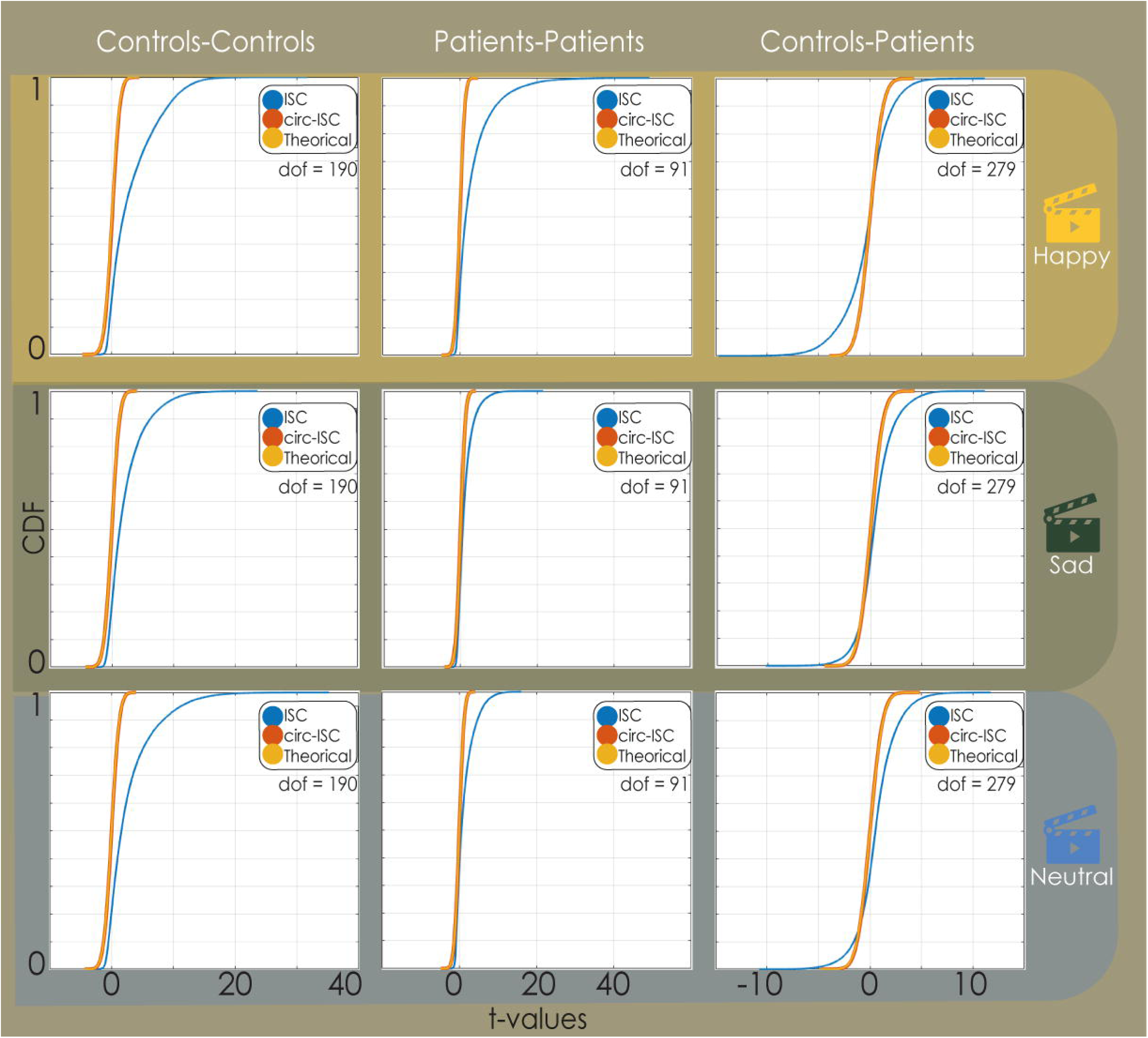

## Footnotes

1 Antipsychotic medication included Risperidona, Quetiapina, Clozapina, Lapenax, and Alental; Antidepressants included Velafaxina, and Paroxetina; Anxiolytics included Lorazepam, Clonazepam, Nozhom, and Logrea

## Notes

### Competing Interest Statement

The authors have declared no competing interest.

