## Supplementary for "Disrupted Emotional Neural Synchrony in Schizophrenia Revealed by Intersubject Correlation of Naturalistic fMRI"

**Running title:** ISC Shows Affective Processing Deficits in Schizophrenia

**Keywords:** Schizophrenia, Intersubject Correlation (ISC), Naturalistic fMRI

Emotional Processing, Neural Synchrony, Social Cognition.

| **Happy** | | | | | |
| --- | --- | --- | --- | --- | --- |
| **Controls** | | | **Schizophrenia** | | |
| **AAL region** | **t-value** | **MNI (x, y, z)** | **AAL region** | **t-value** | **MNI (x, y, z)** |
| Calcarine sulcus | 16.39 | (9, -78, 4) | Calcarine sulcus | 28.12 | (15, -79, 4) |
| Lingual gyrus | 19.73 | (-18, -42, 5) | Lingual gyrus | 31.13 | (17, -45, 1) |
| Middle temporal gyrus | 17.05 | (-59, -10, -5) | Caudate | 34.14 | (-4, 13, -3) |
| Superior temporal gyrus | 15.01 | (67, -8, 8) | Middle occipital gyrus | 18.71 | (-49, -67, 6) |
| Inferior temporal gyrus | 18.24 | (47, -69, -8) | Inferior temporal gyrus | 16.91 | (47, -63, -2) |
| Caudate | 17.66 | (-11, 17, 10) | Thalamus | 21.79 | (-13, -31, 7) |
| **Sad** | | | | | |
| Inferior occipital gyrus | 25.28 | (47, -72, -12) | Parahippocampal gyrus | 13.21 | (-22,-7, -29) |
| Lingual gyrus | 20.46 | (-18, -44, -5) | Calcarine sulcus | 8.82 | (-20, -57, 5) |
| Middle temporal gyrus | 17.38 | (-61, -37, 1) | Lingual gyrus | 10.87 | (-20, -44, -5) |
| Fusiform gyrus | 24.50 | (-42, -72, -16) | Thalamus | 11.99 | (15, -27, 6) |
| Amygdala | 17.55 | (-24, -2, -16) | Inferior temporal gyrus | 10.11 | (44, -63, -8) |
| Precuneus | 18.39 | (-22, -50, 4) | Middle temporal gyrus | 9.45 | (62, -24, -3) |
| **Neutral** | | | | | |
| Middle temporal gyrus | 19.38 | (-50, -24, -1) | Inferior occipital gyrus | 15.09 | (43, -76, -13) |
| Superior temporal gyrus | 20.14 | (58, -16, 1) | Middle temporal gyrus | 14.10 | (-62, -31, 1) |
| Lingual gyrus | 14.43 | (-7,-76, 1) | Inferior temporal gyrus | 14.35 | (48, -61, -5) |
| Inferior occipital gyrus | 15.55 | (-41, -72, -10) | Superior frontal gyrus | 13.60 | (-14, 41, 31) |
| Inferior temporal gyrus | 16.97 | (45,-63, -10) | Lingual gyrus | 9.95 | (-28, -88, -12) |
| Fusiform gyrus | 17.22 | (-38, -72, -13) | Superior temporal gyrus | 7.33 | (57, 37, 12) |

**Supplementary Table 1:** AAL regions, t-values and MNI coordinates corresponding to the local ISC peaks for the three conditions (happy, sad, neutral) and two groups (controls and schizophrenia).

| **ISC (Happy vs. neutral)** | | | | | |
| --- | --- | --- | --- | --- | --- |
| **Controls** | | | **Schizophrenia** | | |
| **AAL region** | **t-value** | **MNI (x, y, z)** | **AAL region** | **t-value** | **MNI (x, y, z)** |
| Insula | 8.88 | (-42, -2, 3) | Calcarine sulcus | 22.87 | (18, -46, 5) |
| Hippocampus | 10.04 | (21, -5, -17) | Lingual gyrus | 19.14 | (8, -66, 2) |
| Amygdala | 9.16 | (-22, -6, -16) | Caudate | 21.37 | (-6, 15, 0) |
| Inferior temporal gyrus | 14.22 | (-37, 4, -34) | Thalamus | 16.62 | (-13, -30, 8) |
| Superior temporal gyrus | -9.88 | (58, -18, 1) | Middle temporal gyrus | 10.86 | (-54, -61, 8) |
| Supplementary motor area | -7.04 | (-9, -8, 63) | Postcentral gyrus | 11.85 | (65, -10, 23) |
| **ISC (Sad vs. neutral)** | | | | | |
| Insula | 6.40 | (-35, -14, 21) | Lingual gyrus | 7.99 | (-17, -44, -3) |
| Lingual gyrus | 11.23 | (-16, -48, 1) | Fusiform gyrus | 7.75 | (26, -44, -12) |
| Thalamus | 9.42 | (-7, -12, 1) | Lingual gyrus | 10.87 | (-20, -44, -5) |
| Amygdala | 13.34 | (-24,-3, -11) | Thalamus | 9.06 | (14, -27, 6) |
| Superior temporal gyrus | -5.08 | (47, -11, -9) | Anterior cingulate cortex | -6.76 | (14, 46, 8) |
| Middle temporal gyrus | -5.17 | (59, -36, -3) | Middle temporal gyrus | -7.73 | (52, -35, 4) |

**Supplementary Table 2:** AAL regions, t-values and MNI coordinates corresponding to the local peaks in the comparisons happy vs. neutral and sad vs. neutral within the controls and schizophrenia groups. Positive / negative t-values indicate greater / less ISC than neutral.

| **ISC (Happy)** | | | | | |
| --- | --- | --- | --- | --- | --- |
| **Controls > schizophrenia** | | | **Controls < schizophrenia** | | |
| **AAL region** | **t-value** | **MNI (x, y, z)** | **AAL region** | **t-value** | **MNI (x, y, z)** |
| Insula | 6.49 | (-41, 1, 6) | Superior frontal gyrus | 7.39 | (13, 37, 38) |
| Middle temporal gyrus | 5.96 | (-55, -31, 6) | Orbitofrontal cortex | 6.16 | (-29, 45, -11) |
| Precentral gyrus | 7.75 | (-46, 4, 53) | Fusiform gyrus | 6.86 | (30, -40, -11) |
| Amygdala | 7.35 | (-24, -3, -13) | Angular gyrus | 10.23 | (-48, -64, 38) |
| Middle temporal gyrus | 5.80 | (-60, -32, 4) | Middle temporal gyrus | 9.32 | (-53, -64, 4) |
| Inferior temporal gyrus | 9.86 | (-39, 9, -36 ) | Precentral gyrus | 8.43 | (-32, -9, 48) |
| **ISC (Sad)** | | | | | |
| Insula | 8.56 | (-36, -18, 14) | Supramarginal gyrus | 6.42 | (61, -29, 42) |
| Superior temporal gyrus | 7.73 | (-42, -33, 14) | Superior frontal gyrus | 8.20 | (-10, 23, 47) |
| Middle temporal gyrus | 7.04 | (54, -45, 4) | Inferior temporal gyrus | 4.36 | (59, -35, -21) |
| Precentral gyrus | 8.90 | (-57, 13, 31) | Middle frontal gyrus | 7.08 | (35, 43, 6) |
| Amygdala | 7.70 | (-22, -6, -12) | Middle occipital gyrus | 5.67 | (47, -83, 6) |
| Fusiform gyrus | 8.55 | (-41, -71, -16) | Supplementary motor area | 6.60 | (-9, 21, 54) |
| **ISC (Neutral)** | | | | | |
| Lingual gyrus | 8.26 | (-9, -71, -3) | Superior frontal gyrus | 6.52 | (-16, 49, 34) |
| Superior temporal gyrus | 6.44 | (58, -18, 2) | Inferior frontal gyrus | 4.64 | (-50, 25, 0) |
| Calcarine sulcus | 5.09 | (17, -71, 12) | Anterior cingulate cortex | 7.27 | (13, 45, 22) |
| Middle temporal gyrus | 4.62 | (60, -44, 12) | Inferior frontal gyrus | 6.23 | (34, 28, 27) |
| Thalamus | 4.56 | (-11, -27, 4) | Superior frontal gyrus | 6.13 | (15, 48, 27) |
| Parahippocampal gyrus | 5.42 | (-20, -37, -9) | Orbitrofrontal cortex | 5.46 | (-10, 58, 0) |

**Supplementary Table 3:** AAL regions, t-values and MNI coordinates corresponding to the local peaks in the comparison controls vs. schizophrenia for the conditions happy, sad and neutral.

| **Happy - neutral** | | | | | |
| --- | --- | --- | --- | --- | --- |
| **Controls > schizophrenia** | | | **Controls < schizophrenia** | | |
| **AAL region** | **t-value** | **MNI (x, y, z)** | **AAL region** | **t-value** | **MNI (x, y, z)** |
| Insula | 5.62 | (-44, 0,4) | Parahippocampal gyrus | 9.53 | (27, -39, -6) |
| Inferior temporal gyrus | 7.74 | (-37, 8 -36) | Lingual gyrus | 8.11 | (-21, -62, -6) |
| Amygdala | 6.43 | (-24, -3, -12) | Inferior occipital gyrus | 7.03 | (-28, -79, -3) |
| Middle temporal gyrus | 4.93 | (-61, -33, -5) | Calcarine sulcus | 8.78 | (15, -77, 5) |
| Superior frontal gyrus | 6.37 | (-9, 57, 20) | Thalamus | 7.42 | (9, -28, 5) |
| Putamen | 4.72 | (-24, 2, -6) | Cuneus | 8.07 | (9, -81, 23) |
| **Sad - neutral** | | | | | |
| Insula | 5.23 | (-34, -18, 16) | Calcarine sulcus | 5.15 | (-6, -66, 11) |
| Amygdala | 5.75 | (-22, -5, -16) | Middle frontal gyrus | 5.65 | (34, 56, 1) |
| Amygdala | 5.98 | (22, -3, -16) | Thalamus | 4.13 | (-13, -24, 10) |
| Middle temporal gyrus | 6.68 | (48, -70, 2) | Middle occipital gyrus | 5.63 | (37, -81, 28) |
| Fusiform gyrus | 7.92 | (43, -73, -16) | Superior frontal gyrus | 4.92 | (9, 24, 37) |
| Inferior frontal cortex | 5.94 | (-58, 13, 28) | Precentral gyrus | 4.29 | (45, -9, 60) |

**Supplementary Table 4:** AAL regions, t-values and MNI coordinates corresponding to the local peaks in the comparison controls vs. schizophrenia for the conditions happy and sad with neutral subtracted as a baseline.

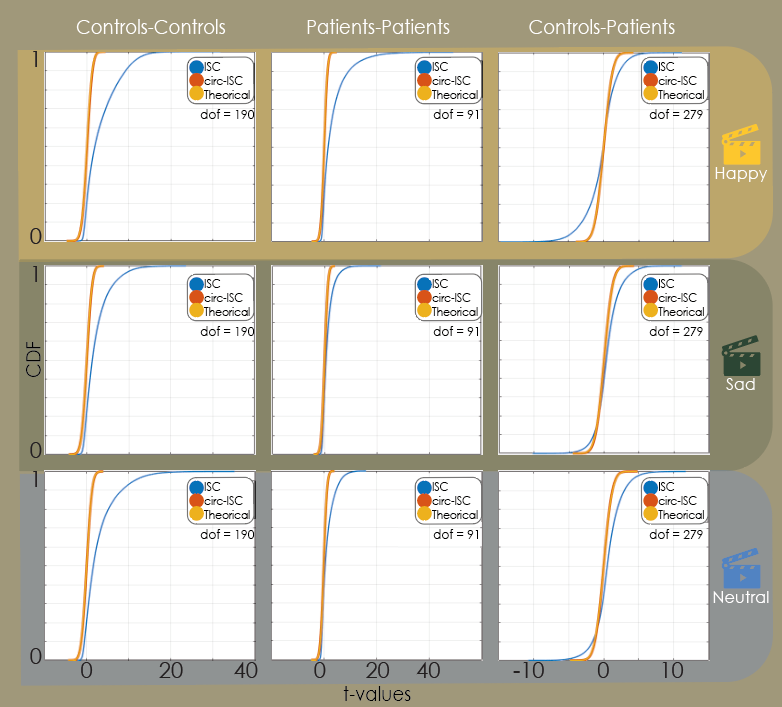

**Supplementary Figure 1. Cumulative distribution functions (CDF) of t-values from the Intersubject Correlation (ISC) analyses.** Columns show within-controls, within-patients, and between-groups comparisons; rows represent the three emotional conditions (Happy, Sad, Neutral). Curves depict observed ISC (blue), circularly shifted ISC (orange), and the theoretical null distribution (gray). Degrees of freedom (dof) correspond to the number of subject pairs per contrast (controls = 190, patients = 91, between-groups = 279).
